# Growth rate trades off with enzymatic investment in soil filamentous fungi

**DOI:** 10.1101/360511

**Authors:** Weishuang Zheng, Anika Lehmann, Masahiro Ryo, Kriszta Kezia Vályi, Matthias C. Rillig

## Abstract

Saprobic soil fungi drive many important ecosystem processes, including decomposition, and many of their effects are related to growth rate and enzymatic ability. In mycology, there has long been the implicit assumption of trade-off between growth and enzymatic investment, which we here test. Using a set of 31 filamentous fungi isolated from the same ecosystem, we measured growth rate (as colony radial extension) and enzymatic repertoire (activities of four enzymes: laccase, cellobiohydrolase, leucine aminopeptidase and acid phosphatase). Our results support the existence of a trade-off, however only for the enzymes representing a larger metabolic cost (laccase and cellobiohydrolase). Our study offers new insights into functional complementarity within the soil fungal community in a number of ecosystem processes, and experimentally supports an enzymatic investment/ growth rate tradeoff in explaining phenomena including substrate succession.

## Introduction

Soil saprobic fungi drive essential ecosystem processes including decomposition and soil aggregation. In order to understand contributions of fungi to such processes it is helpful to adopt a functional traits approach (Lehmann and Rillig, 2015). Two traits are expected to be of great importance for describing essential functions of fungi in soil: growth rate, because it denotes how fast and to what extent fungi encounter their substrate (Crowther *et al*., 2014); and enzymatic repertoire, since it describes the ability to break down and utilize various substrates (Baldrian *et al*., 2011; Eichlerová *et al*., 2015).

Traditional concepts in mycology have long assumed a fundamental tradeoff between growth rate and enzymatic abilities. For example, substrate successions have been described as a shift in fungal strategies where the ability to grow fast and the ability to use more recalcitrant carbon compounds change over time (Kendrick and Burges, 1962; Frankland, 1998). For ectomycorrhizal fungi, initial experimental evidence (comparing three species; Moeller and Peay, 2016) and observations (Agerer, 2001) have suggested a tradeoff between investment in enzymes and exploration distance or competitiveness. Nevertheless, the existence of such a tradeoff has never been directly experimentally addressed in soil filamentous fungi.

We here use a set of 31 fungi, isolated from the same soil and representing three fungal phyla (Ascomycota, Basidiomycota and Mucoromycotina), and ask in dedicated experiments if and how growth rate and enzymatic repertoire is linked. Employing a multiple threshold approach (Byrnes *et al*., 2014; Lefcheck *et al*., 2015), we additionally explore joint patterns of enzyme activities in relation to growth rate.

## Materials and methods

The filamentous saprobic fungi used in our experiments were isolated from soil samples collected in one ecosystem, a grassland at Oderhänge Mallnow (Germany, 52°27.778’ N, 14°29.349’ E). The 31 isolates belong to the Ascomycota, Basidiomycota and Mucoromycotina (Fig. 1; Andrade-Linares *et al*., 2016). Fungi were grown in Petri dishes with potato dextrose agar: this is a rich medium in which all fungi can growth without overt nutrient limitation, and fungal strains can also invest in enzyme production (Allison *et al*., 2014). Furthermore, we did not use lignin or other substances to trigger activity of specific enzymes. We placed sterilized cellophane membranes (#165-0963, Bio-RAD, USA) on the agar surface to facilitate extraction of media-free mycelia (Zheng *et al*., 2014). We grew fungi from mycelium plugs at room temperature (25˚C) until colonies were past half of their linear growth phase. Six replicates were prepared for each strain resulting in 186 experimental units.

**Figure 1.**
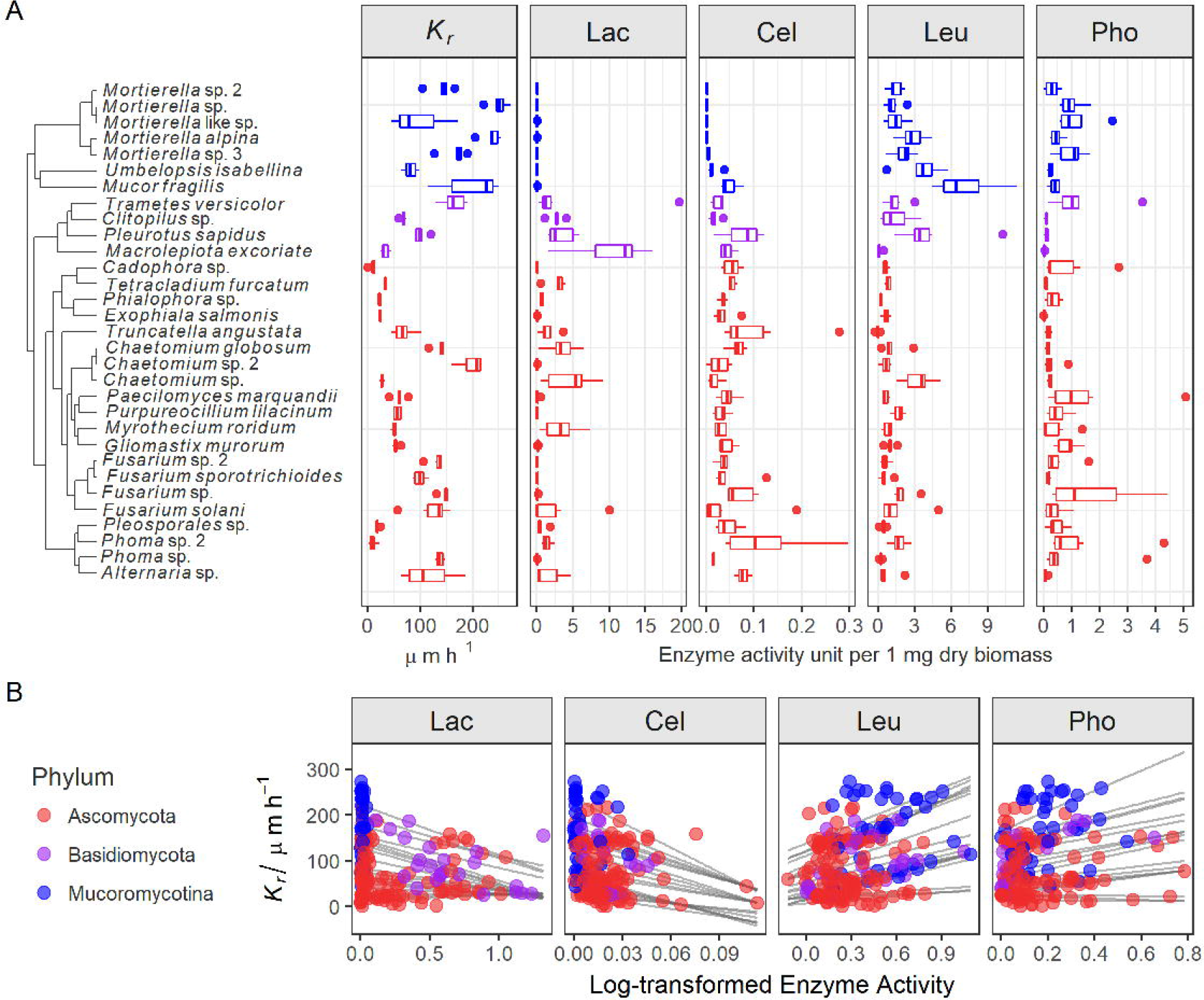
Traits of 31 saprobic fungi and the interaction between colony radial growth rate (*K_r_*) and activities of laccase (Lac), cellobiohydrolase (Cel), leucine aminopeptidase (Leu) and acid phosphatase (Pho). **Panel A**: phylogenetic distribution of *K_r_* and the enzymes (6 replicates). **Panel B**: quantile regression between *K_r_* and log-transformed enzyme activity data (n=186). For Lac, the regression starts to be significant from the 23% quantile, resulting in a triangle-shape. This is caused by the lack of Lac in Mucoromycotina species. The R^2^ of regression on the mean for each enzyme are 0.08 (Lac), 0.07 (Cel), 0.08 (Leu), 0.03 (Pho). Details of the statistics are in Table S3.

### Colony radial growth rate (*K_r_*)

All colonies were imaged (Epson Perfection V700) at two time points. In each picture, the radius was measured in triplicate with Image-J (1.49 v23; Schneider et al., 2012), and subsequently averaged to calculate the growth rate, following the equation: 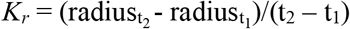.

### Enzymes

We profiled fungal enzymatic activities of laccase (Lac; lignin degradation), cellobiohydrolase (Cel; cellulose degradation), leucine aminopeptidase (Leu; catalyzes the hydrolysis of peptides) and acid phosphatase (Pho; releases free attached phosphoryl groups) by a microplate photometric method following a modified protocol by Courty *et al*. (2006). We cut small pieces of mycelium (3-5 mm^2^) from the colony’s peripheral zone, weighed them, stored at 4°C and processed them within 24 hours.

The relationship between *K_r_* and enzyme activities were tested (i) considering single enzymes and (ii) as a multifunctional property. First, we applied quantile regression with the package “quantreg” (https://github.com/cran/quantreg) on log transformed enzyme activities and *K_r_* (n = 186). Second, we used the first principal component (PC1), extracted from principal component analysis (PCA) on 4 enzyme activities, to represent the enzyme profile, and regressed it with *K_r_* (n = 31). Last, we applied to our question a multiple threshold approach (Byrnes *et al*., 2014) exploring at threshold values from 0.05 to 0.99 the relationships between the number of activated enzymes and *K_r_*. For this, the values of Lac and Cel were reflected. All analyses were conducted in R (3.4.1; R Core Team, 2014). For detailed description of the protocols and statistical procedures see Supplementary Information.

## Results and Discussion

We found high variability in traits across the tested 31 fungi covering 11 orders from three different (sub-)phyla (Fig. 1A). The trait *K_r_* varied from 9.2 to 250.2 µm h^-1^ with the fastest growing strains found in Mucoromycotina. The enzyme profiles showed that Lac activity was much higher in Basidiomycota than in Ascomycota, while it was absent in Mucoromycotina. Mucoromycotina had higher Leu compared to the other strains. Within Mucoromycotina, Mortierellales strains did not produce Cel. All strains had Pho activity. These results are broadly consistent with the known enzymatic features of fungal groups (Baldrian *et al*., 2011; Courty *et al*., 2006; Eichlerová *et al*., 2015).

We found clear evidence of a tradeoff in enzyme activity and growth rate; however, not for all enzymes. Quantile regression analyses of the relationship between *K_r_* and the individual enzymes revealed that strains with higher Lac and/or Cel activities grew more slowly, while those with high Pho and/or Leu activities grew faster (Fig.1B, Table S3). Since we cultivated strains in nutrient-sufficient media rich in starch and dextrose, Lac and Cel were not necessary for growth. We interpret the observed link between growth rate and enzyme activities as an aspect of intrinsic strategy of the fungi, rather than a reaction to their cultivation conditions. Our data support the idea that fast-growing fungi invest in enzymes targeting easily accessed rather than recalcitrant substrates, and vice versa. This suggests also that Lac and Cel enzymes come at a higher metabolic cost to the mycelium, leaving less for allocation to growth. A slightly different perspective with the same outcome was offered by Agerer (2001) based on observations on ectomycorrhizal fungi: those fungi that cannot produce phenoloxidases to decompose lignin will have to explore long-distance for easily decomposed carbon resources, and vice versa.

Under natural conditions, fungi tend to produce a cocktail of enzymes to acquire resources, hence we also explored the impact of the combined effect of the four enzymes on *K_r_*. First, we revealed that compared to single enzymes, the overall enzyme activity (reflected in PC1) has a stronger effect on *K_r_* (Fig. 2A), especially for Basidiomycota strains (purple dots; R^2^ = 0.74, F_1,2_ = 5.805; p = 0.14). This further supports the hypothesis of a tradeoff between growth and enzymatic repertoire, differentiated to specific enzymes. Note that loading of PC1 for Lac and Cel (+) is opposite from that of Pho and Leu (–). Because of these opposite relationships of ‘cheap’ vs. ‘costly’ enzymes with *K_r_*, in the multiple threshold approach, we reflected Lac and Cel.

**Fig. 2.**
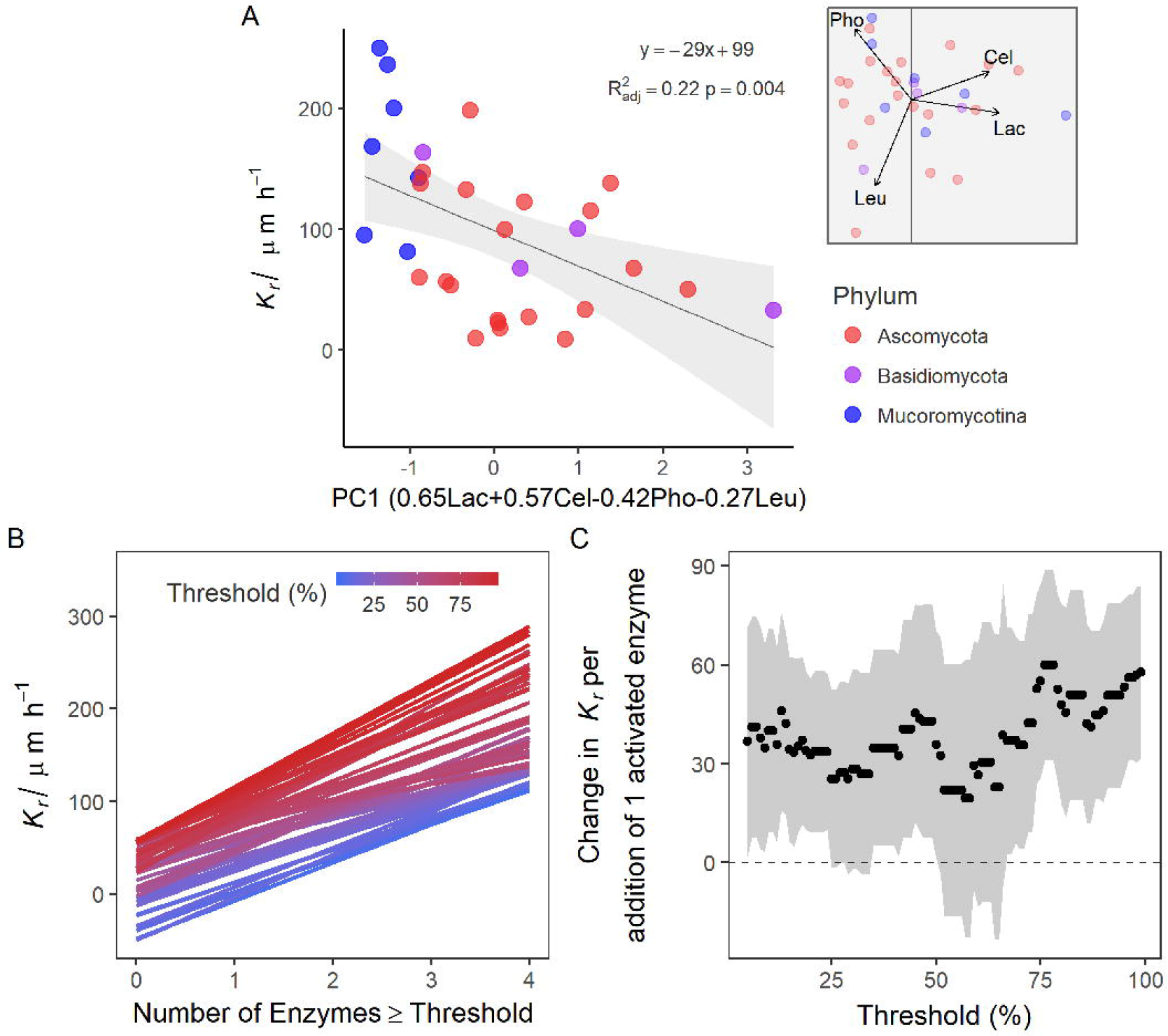
Impact of simultaneous effects of the four enzymes on colony radial growth rate (*K_r_*) of saprobic fungi. **Panel A**: the overall effect of enzymes represented by PC1 on *K_r_*. The PCA results are shown in the small insert. PC1 loads positively on Lac and Cel, and negatively on Leu and Pho. The loadings are shown in the x-axis label. **Panel B and C**: multiple thresholds analysis on *K_r_* and enzyme activities. **Panel B** shows relationship between *K_r_* and the number of enzymes activated above a threshold for multiple different threshold values. Color bars within panel B indicate different thresholds (blue = 0 %, red = 100 %). **Panel C** shows the corresponding relationship between threshold value and slope of the regression lines in panel B. Points are the estimated slope values and shading indicates confidence intervals (CIs). At the whole range of thresholds, except 25–34% and 51–66%, the CIs of slopes does not overlap with zero. The maximal effect of multiple enzymes occurs at a threshold value of 76% with the slope of 59.9 (details of the metrics are in Tables S3).

The multiple threshold approach confirms that the enzymes had strong ‘multifunctionality’ effects on *K_r_*, i.e., enzymes have combined relationships with growth rate (Fig. 2B and C). These effects occurred not only at high thresholds (66–99%), but also at low threshold values, indicating a high baseline of multifunctionality. To have a high *K_r_*, a fungus has to invest more into relatively ‘cheap’ enzymes (here: Pho and Leu) and less in ‘costly’ enzymes (here: Lac and Cel, note that these were reflected); and vice versa for low growth rates. Even though we measured only a limited number of enzymes, our result is the first empirical evidence showing ‘multifunctionality’-effects of enzymes on *K_r_*.

In conclusion, we show that, for saprobic fungi across several phyla, growth rate is related to multiple enzyme activities, and depending on whether enzymes are targeting recalcitrant substrates or not, enzymes are linked positively or negatively with fungal growth rate. Our analysis also highlights the combined effect of the entire enzymatic repertoire, to the extent measured, rather than a key enzyme as responsible for this pattern. Our results thus offer new insights into potential complementarity in the soil fungal community in a number of ecosystem processes, and assert an enzymatic investment/ growth rate tradeoff in explaining phenomena including substrate succession.

## Acknowledgements

MCR acknowledges funding from Deutsche Forschungsgemeinschaft (RI 1815/16-1) and from the ERC Advanced Grant ‘Gradual Change’ (694368) for this work. WZ acknowledges support from the China Scholarship Council.

## Conflicts of interest

The authors declare no conflicts of interest.

## Notes

Conflict of interest: The authors declare no conflicts of interest.

## References

Agerer R. (2001). Exploration types of ectomycorrhizae A proposal to classify ectomycorrhizal mycelial systems according to their patterns of differentiation and putative ecological importance. Mycorrhiza 11: 107–114.

Allison SD, Chacon SS, German DP. (2014). Substrate concentration constraints on microbial decomposition. Soil Biol Biochem 79: 43–49.

Andrade-Linares DR, Veresoglou SD, Rillig MC. (2016). Temperature priming and memory in soil filamentous fungi. Fungal Ecol 21: 10–15.

Baldrian P, Voříšková J, Dobiášová P, Merhautová V, Lisá L, Valášková V. (2011). Production of extracellular enzymes and degradation of biopolymers by saprotrophic microfungi from the upper layers of forest soil. Plant Soil 338: 111–125.

Byrnes JEK, Gamfeldt L, Isbell F, Lefcheck JS, Griffin JN, Hector A, et al. (2014). Investigating the relationship between biodiversity and ecosystem multifunctionality: Challenges and solutions. Methods Ecol Evol 5: 111–124.

Courty PE, Pouysegur R, Buée M, Garbaye J. (2006). Laccase and phosphatase activities of the dominant ectomycorrhizal types in a lowland oak forest. Soil Biol Biochem 38: 1219–1222.

Crowther TW, Maynard DS, Crowther TR, Peccia J, Smith JR, Bradford MA. (2014). Untangling the fungal niche: The trait-based approach. Front Microbiol 5: 1–12.

Eichlerová I, Homolka L, Žif čáková L, Lisá L, Dobiášová P, Baldrian P. (2015). Enzymatic systems involved in decomposition reflects the ecology and taxonomy of saprotrophic fungi. Fungal Ecol 13: 10–22.

Frankland JC. (1998). Fungal succession — unravelling the unpredictable. Mycol Res 102: 1–15.

Kendrick W, Burges A. (1962). Biological aspects of the decay of *Pinus sylvestris* leaf litter. Nov Hedwigia 4: 313–342.

Lefcheck JS, Byrnes JEK, Isbell F, Gamfeldt L, Griffin JN, Eisenhauer N, et al. (2015). Biodiversity enhances ecosystem multifunctionality across trophic levels and habitats. Nat Commun 6: 1–7.

Lehmann A, Rillig MC. (2015). Understanding mechanisms of soil biota involvement in soil aggregation: A way forward with saprobic fungi? Soil Biol Biochem 88: 298–302.

Moeller H V., Peay KG. (2016). Competition-function tradeoffs in ectomycorrhizal fungi. PeerJ 4: e2270.

R Core Team. (2014). R: A language and environment for statistical computing. http://www.rproject.org/.

Schneider CA, Rasband WS, Eliceiri KW. (2012). NIH Image to ImageJ: 25 years of image analysis. Nat Methods 9: 671–675.

Zheng W, Morris EK, Rillig MC. (2014). Ectomycorrhizal fungi in association with *Pinus sylvestris* seedlings promote soil aggregation and soil water repellency. Soil Biol Biochem 78: 326–331.

